# XGMix: Local-Ancestry Inference with Stacked XGBoost

**DOI:** 10.1101/2020.04.21.053876

**Authors:** Arvind Kumar, Daniel Mas Montserrat, Carlos Bustamante, Alexander Ioannidis

**Author notes:** Work conducted during an internship at Stanford University (Daniel Mas Montserrat).

## Abstract

Genomic medicine promises increased resolution for accurate diagnosis, for personalized treatment, and for identification of population-wide health burdens at rapidly decreasing cost (with a genotype now cheaper than an MRI and dropping). The benefits of this emerging form of affordable, data-driven medicine will accrue predominantly to those populations whose genetic associations have been mapped, so it is of increasing concern that over 80% of such genome-wide association studies (GWAS) have been conducted solely within individuals of European ancestry [1]. The severe under-representation of the majority of the world’s populations in genetic association studies stems in part from an addressable algorithmic weakness: lack of simple, accurate, and easily trained methods for identifying and annotating ancestry along the genome (local ancestry). Here we present such a method (XGMix) based on gradient boosted trees, which, while being accurate, is also simple to use, and fast to train, taking minutes on consumer-level laptops.

## 1 Introduction

Genome-wide association studies (GWAS) are based upon correlations between observed (genotyped) sites and causal sites in their vicinity. (The majority of DNA sites, including most causal sites, are not amongst the small number of genotyped sites.) However, due to the independence of genetic drift in separated populations [2, 3], long-separated populations exhibit markedly different correlations between neighboring variants along the genome. Thus, association models trained on one population have inherent biases, and are several times less accurate when applied to another population [4]. Indeed, in certain cases the same variant may have a significant, opposite, association with a disease depending on the population’s ancestry [5]. Thus, it is critical that genetic associations be mapped in all populations. However, this is not happening [1][6]; instead, a new divide is rising in healthcare between the small number of populations (predominantly European) that have been included in genetic association studies, and the majority of the world’s populations, which have not been studied and cannot take advantage of these precise personalized predictions. This gap is emerging not due to researchers lacking access to diverse patients to study, but rather because most association studies deliberately exclude diversity. Such studies focus on single ancestry, preferably homogeneous, cohorts to increase statistical power, thereby avoiding the issue of differing associations along the genome in differing ancestries. This approach necessarily excludes all admixed groups, for instance African American or Hispanic populations, who encompass more than one ancestry, and also avoids populations that contain too much genetic variation, or too many diverse sub-populations, as is common within Africa.

Fortunately, an algorithmic solution exists to the problem of complex genomic covariance structures in admixed and diverse populations: local ancestry inference (LAI) (figure 1). By using pattern-matching algorithms along the genome, local ancestry inference can annotate each region of the genome with an ancestry label. This permits association studies to be run using an indicator label (ancestry at each genomic position) to de-convolve the effects of ancestry on the correlation structure of neighboring genomic variants [7]. It also allows accurate personalized genetic predictions (polygenic risk scores) to be based on the ancestry of the genomic region in which an associated variant is located. Unfortunately, current local ancestry inference methods are both difficult to use and computationally expensive [8, 9].

**Figure 1:**
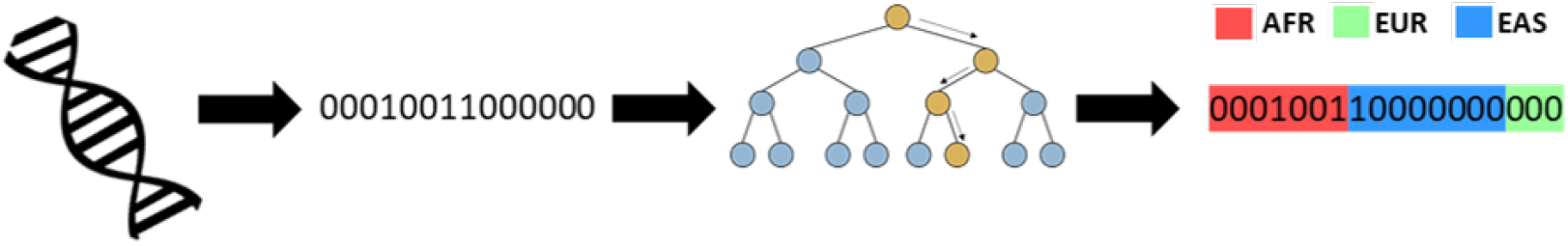
Local-ancestry inference problem. Each chromosome is genotyped and represented as a binary string of SNP variants. A LAI method (such as XGMix) identifies the ancestry for each SNP along the genome.

Currently, most pattern recognition problems are tackled via deep learning-based methods. Indeed, neural networks have proven to be both accurate and efficient, in many cases surpassing the performance of methods based on hand-crafted features, and even, human performance. Typically, such neural networks require GPUs for their training and inference. This can be restrictive, as GPUs are expensive, and alternatives such as online compute services tend to be costly, if used extensively. Community diagnostic labs and regional medical facilities in developing countries could find access to GPU computing, not to mention access to compute support staff, prohibitively expensive, and could find cloud connectivity to be unreliable. Such computational resource limits create a need for methods that can be trained and deployed without technical support and with low computing requirements, such as consumer laptops.

In this work, we present a method based on XGBoost systems [10]. XGBoost is a gradient tree boosting method that has provided state-of-the-art results in many tasks while being scalable and computationally efficient. Because XGBoost is available in many programming languages, frameworks, and distributed computing environments, an XGBoost-based method can easily be integrated into other biomedical processing applications. Its parallel nature allows for it to be trained fast on multi-core laptops without the need for GPUs.

The presented method, XGMix, estimates an ancestry label within windowed sections of haploid human chromosome sequences, providing results that are competitive with previous state-of-the-art LAI methods. Additionally, our architecture optionally permits approaching local-ancestry inference as a regression problem by estimating the coordinates (latitude and longitude) representing the geographic source location for the ancestry of each window of each chromosome. Rather than inferring a small number of discrete ancestry classes, by inferring the ancestral source coordinates of each segment of the chromosome, we can accommodate diverse and continuous ancestry variation, as is common across most regions of the world outside the post-colonial Americas.

## 2 Related Work

Several methods for local-ancestry inference have been published. SABER [11], HAPAA [12] and HAPMIX [13] estimate local-ancestry for genomic sequences using Hidden Markov Models. The accuracy and efficiency of these methods are surpassed by LAMP [8], a method that applies probability maximization within a sliding window. SVMs [14] and Random Forests with a conditional random field (RFMix) [9] have also been explored for increased accuracy, with the latter currently considered to be state-of-the-art. Recently, LAI-Net [15], a local-ancestry estimation method based on a neural network, was demonstrated to provide competitive results with RFMix, while a class-conditional variational autoencoder generative adversarial network (C-VAE-GAN) has been used to create synthetic single-ancestry and admixed individuals from a set of desired ancestries [16].

Besides Random Forests, which have proven so successful in the local ancestry problem, other classifiers and regressors based on decision trees (CART) exist, including AdaBoost [17], based on adaptive boosting, and XGBoost [10], based on gradient boosting. Gradient tree boosting methods, also known as gradient boosting machines (GBM) or gradient boosted regression tree (GBRT), have proven to be more accurate than other tree-based approaches and have provided state-of-the-art results in many fields [10, 18].

## 3 Admixture Dataset

We used a dataset composed of full genome sequences from admixed individuals generated from real worldwide populations from the 1000 genomes project [19]. We chose single-population-ancestry individuals from Africa (AFR), East Asia (EAS), and Europe (EUR) selecting one population from each of these continents: 108 Yoruba from Ibadan, Nigeria (YRI), 103 Han Chinese from Beijing, China (CHB), and 107 Spanish individuals (IBS). Using these real individuals’ sequences we simulated descendants with admixed ancestry using Wright-Fisher forward simulation over a series of generations using standard human recombination rate parameters [9]. Simulated admixed individuals must be used for training, rather than real admixed individuals, so that we know the ancestry switch points along the chromosome. From these 318 real single-ancestry individuals, we selected 258 at random to generate 600 admixed individuals for training. Thirty individuals (ten from each group) were selected to generate 300 admixed individuals for validation and the remaining thirty individuals (ten from each group) were used to generate 300 admixed individuals for testing. The training set, composed of 600 individuals, consisted of 6 groups of 100 individuals generated by 2, 8, 12, 32, 48 and 64 generations of admixture. The validation and testing set, composed of 300 individuals each, consisted of 3 groups of 100 individuals generated by 4, 16, and 24 generations of admixture. As the number of generations increases following initial admixture, descendants have more frequent ancestry switches along the genome, thus smaller fragments of their genome are inherited together from the same ancestry, leading to more challenging inference.

Additionally, we generated a dataset with genetically similar populations to perform an evaluation of the method’s performance when faced with a more challenging discriminative task. We simulated 400 admixed individuals using real human genomes from individuals sampled across Asia: 182 Han Chinese (CHB and CHS), 83 Chinese Dai (CDX), 89 Vietnamese Kinh (KHV), 94 Japanese (JPT), 93 Gujarati Indians (GIH), 86 Pakistani Punjabi (PJL), 76 Bangladeshi Bengali (BEB), 92 Sri Lankan Tamil (STU) and 92 Indian Telugu (ITU). We generated 200 admixed individuals for training using a different set of 100 real Asian individuals, ten from each population. The remainder of the real individuals were used to generate 200 simulated admixed individuals for training. Both training and testing individuals were generated after two and four generations. Since local-ancestry inference methods must accurately estimate the ancestry from individuals regardless of their admixture histories (different generation times since admixture), it is important to train and evaluate the method with admixed individuals simulated over a wide range of generations.

## 4 Proposed Method

The proposed method (XGMix) consists of a set of stacked XGBoost trees. A first level of parallel XGBoosts performs an initial estimation of the ancestry. A unique XGBoost then passes the initial estimates through a sliding window and outputs the final ancestry estimates. We show how XGMix can perform local-ancestry inference both via classification and via geographical coordinate regression. When geographical coordinate regression is used, a category can be assigned post hoc by matching the coordinate with the closest ground-truth coordinate. Figure 2 presents the structure of XGMix based on ancestry classification.

**Figure 2:**
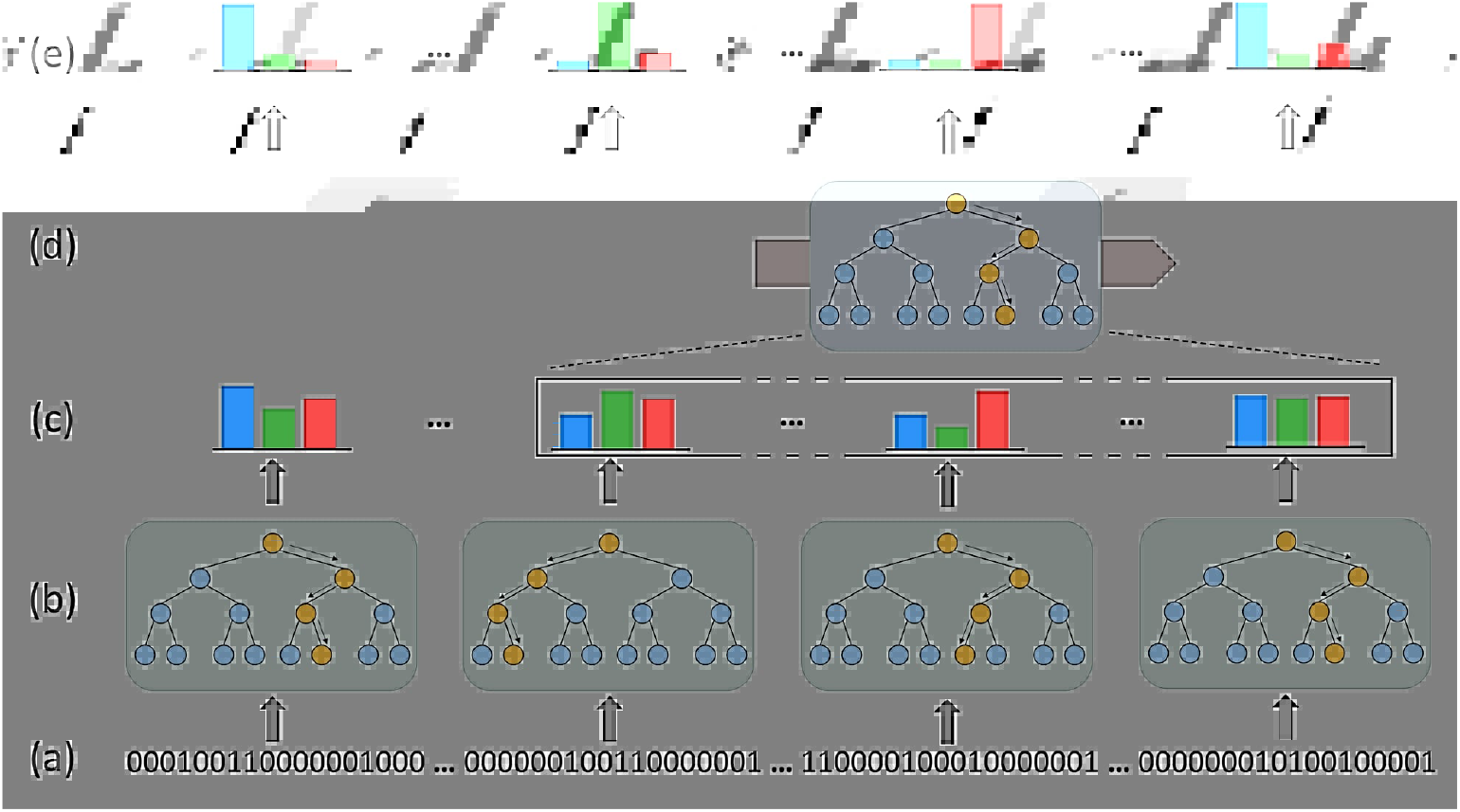
Diagram of the XGMix method: (a) Input haploid sequences are divided into non-overlapping fixed-size windows. (b) Sets of XGBoost systems applied to each window. (c) Initial probability estimates for each window for each ancestry class (colored bars). (d) Sliding XGBoost system that locally smooths initial estimates. (e) Final probability estimates for each ancestry class.

The first layer of gradient boosting trees takes as input the haploid sequences of SNPs (maternal and paternal sequences are treated independently). The binary nature of the chromosome makes tree-based approaches a good fit. Because the data is not translation invariant (an identical sequence of 0s and 1s might indicate a different ancestry depending its location in the genome) a different tree is applied at each windowed section of the input sequence. We use a window size of 500 and within each window we apply an XGBoost system composed of 100 trees with a depth of 4 levels and a learning rate of 0.1. (These parameters of our system were selected with a validation set). Each tree outputs the probability of the window belonging to a predefined ancestry (in the classification-based XGMix), or the geographical coordinate indicating the likely sampling location (in the coordinate regression-based XGMix).

The second layer includes one sliding gradient boosting tree that smooths the initial estimates. This sliding XGBoost system takes as input the initial probabilities (or coordinates) from the current window and 100 neighboring windows (similar to a convolution operator) and outputs a final estimate. It is composed of 100 trees with a depth of four levels and a learning rate of 0.3. This approach differs from previous smoothing techniques, such as the use of a Conditional Random Field (CRF) by RFMix [9], which requires the user to input parameters, such as the number of generations since the admixing process occurred. By using a sliding window XGBoost can train the smoother in a data-driven way to learn the statistical properties of ancestry switches within chromosome sequences without stipulating the time since admixture, which is generally not known, and may also vary across a population.

While previous LAI methods have framed this problem as a classification task, our regression approach has advantages. A classification task treats each ancestry misclassification equally, even though some ancestries are much more related than others. Thus, LAI methods break when faced with very closely related populations, while a regression formulation can still provide useful estimates. We combine XGMix (coordinate regression) with a Gaussian kernel density estimator [20] to estimate a probabilistic geographical representation that can be useful in downstream applications. Examples are shown in figure 3 and section 5.

## 5 Experimental Results

We used the simulated admixed dataset to train and test XGMix and a validation set to select the hyperparameters: window size, number of trees per windows, levels of trees and learning rates. (While the selection of window size lead to different accuracy results, the number and level of trees, and learning rate, did not have a significant effect on the accuracy on the method.) Table 1 presents the accuracy results for chromosome 20 of XGMix based in classification and regression with and without the smoothing layer as compared with RFMix accuracy on the same task.

**Table 1:** Accuracy of RFMix compared with XGMix with smoothing and without smoothing layer in the validation and testing set.

| Method | Smoothing | Validation | Test |
| --- | --- | --- | --- |
| RFMix [9] | - | 95.19% | 93.00% |
| XGMix (Classification) | No | 78.30% | 78.59% |
|  | Yes | 98.20% | 97.98% |
| XGMix (Coord Regression) | No | 78.30% | 78.34% |
|  | Yes | 96.63% | 96.59% |

Tests suggest that both XGMix based on classification and XGMix based on regression achieve state-of-the-art accuracy with no significant difference between the classification and regression models. The method can be trained within ~30 min in a consumer-level laptop (an operation that need be performed only once), while methods such as RFMix require an order of magnitude longer.

Genotype data can be incomplete due to genotyping errors, or only a subset of SNPs might be available depending on the commercial genotyping array used. Therefore, methods that are robust to missing SNP data are preferred. We evaluate the performance of XGMix when large amounts of data are missing, training and testing XGMix with different percentages of missing input SNPs. While the input sequence is modeled with 0s and 1s, we assign 2s to the missing values, without changing the tree structure. Table 2 presents the accuracy values of XGMix (based on classification) with a different percentage of missing input SNPs.

**Table 2:** Accuracy of XGMix for different percentage of missing input SNPs.

| Missing SNPs | w/out Smoothing | w/ Smoothing |
| --- | --- | --- |
| **0%** | 78.59% | 97.98% |
| **20%** | 74.88% | 96.78% |
| **50%** | 69.20% | 95.00% |
| **80%** | 58.78% | 93.19% |
| **90%** | 44.72% | 84.20% |

The results show XGMix is able to estimate ancestry without a significant loss of accuracy, even when 80% of the input SNPs are missing. This enables the application of the method to cases where cost constraints, such as large patient throughput, necessitate sparser, reduced-cost genotyping.

We perform a qualitative evaluation of the coordinate regression XGMix on the Asian population dataset. While classification-based approaches fail in closely related populations (obtaining ~15% accuracy in this dataset), a coordinate regression-based approach is able to provide meaningful representations of an individual’s ancestry. Figure 3 shows an example of the estimated density map of dual-ancestry admixed individuals using a model trained on all of the Asian populations.

**Figure 3:**
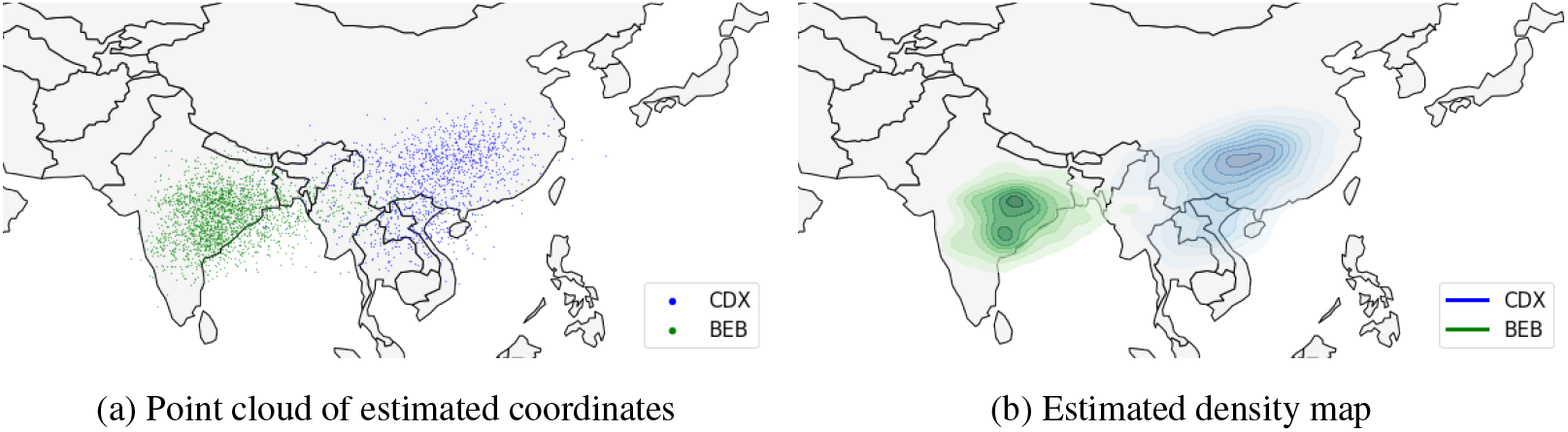
Coordinates and density map with points representing the inferred ancestral locations of each chromosomal segment in a single admixed individual of 70% Bengali (BEB) and 30% Han Chinese (CDX) ancestry, based on the model trained across all Asian populations in 1000 Genomes. The colors of each point correspond to the true ancestral origin of each chromosomal segment.

## 6 Conclusions

Identification and annotation of local ancestry along the genome enables genome-wide association studies to handle admixed and diverse populations. This in turn allows for the expansion of such studies beyond the homogeneous (typically European) populations generally used. Enabling these worldwide studies is crucial, since mapping genetic associations across all of the world’s populations is a prerequisite for extending the rapidly emerging field of affordable, data-driven, personalized genomic healthcare to all. XGMix is extremely accurate, easy to share once trained, simple, computationally inexpensive, and robust to missing data, thus functioning across multiple genotyping platforms, including more economical models.

## A Appendix: Extended Experiments

We performed further experiments to study the effect on accuracy of window size, smoothing size, and testing data generation number. In order to do so, we test XGMix (Classification) on subsets of the validation data that contain simulated admixed individuals of specific generations.

Table 3 shows the effect of smoothing window size on individuals from differing admixture generations. Results show that larger smoothing windows provide better accuracy, except for individuals having large admixture generation values. This is expected, since individuals for whom the admixing process happened many generations ago will have frequent ancestry switches, and therefore segments where the ancestry is constant will be small. In this scenario information from distant windows isn’t useful, and large smoothing window size doesn’t provide accuracy improvement. On the other hand, for individuals where the admixing processes happened recently, the ancestry switch frequency is low, so distant genomic regions are still informative.

**Table 3:** Accuracy (in percent) of various XGMix configurations (classification with window size 500 SNPs) tested on different generations.

| Smoothing<br>Size | Generation Number |  |  |  |  |  |  |  |
| --- | --- | --- | --- | --- | --- | --- | --- | --- |
|  | 2 | 4 | 8 | 12 | 24 | 32 | 48 | 64 |
| **No** | 78.0 | 79.0 | 78.6 | 79.0 | 78.5 | 77.7 | 78.3 | 77.9 |
| **20** | 97.2 | 97.6 | 97.0 | 96.7 | 96.1 | 95.8 | 95.5 | 94.6 |
| **50** | 98.6 | 99.0 | 98.3 | 98.1 | 97.5 | 96.5 | 95.7 | 94.8 |
| **100** | 99.1 | 99.3 | 98.8 | 98.4 | 97.6 | 96.5 | 95.2 | 93.9 |
| **200** | 99.3 | 99.5 | 98.9 | 98.4 | 97.6 | 96.4 | 95.2 | 93.9 |

Table 4 shows the effect of including smoothing in relation to window size. We observe that for small window sizes (500) the accuracy difference with and without smoothing is quite large (~ 17%); however, when using large window sizes (2000), the accuracy difference is lower (~8%). Such behavior is expected, since large window sizes are able to capture relationships between SNPs further away.

**Table 4:** Accuracy (in percent) of various XGMix configurations tested on different generations.

| Window<br>Size | Smoothing<br>Size | Generation Number |  |  |  |  |  |  |  |
| --- | --- | --- | --- | --- | --- | --- | --- | --- | --- |
|  |  | 2 | 4 | 8 | 12 | 24 | 32 | 48 | 64 |
| **500** | **No** | 78.0 | 79.0 | 78.6 | 79.0 | 78.5 | 77.7 | 78.3 | 77.9 |
|  | **50** | 98.6 | 99.0 | 98.3 | 98.1 | 97.5 | 96.5 | 95.7 | 94.8 |
| **1000** | **No** | 84.7 | 86.0 | 85.3 | 85.5 | 85.0 | 84.4 | 84.9 | 84.3 |
|  | **50** | 98.9 | 99.3 | 98.6 | 98.5 | 98.0 | 97.2 | 96.6 | 95.7 |
| **2000** | **No** | 89.5 | 90.0 | 90.0 | 90.0 | 89.5 | 89.1 | 89.6 | 88.8 |
|  | **50** | 98.3 | 98.6 | 98.0 | 97.8 | 97.4 | 96.5 | 96.5 | 95.7 |

A common behavior that can be observed in tables 3 and 4 is an accuracy decrease with increasing number of admixture generations. This is expected, since a higher admixture generation number implies more frequent ancestry switches and thus shorter sequences with constant ancestry. This makes switches more challenging to detect.

Table 5 shows the mean absolute error of XGMix (regression) in both admixed simulated datasets. Although the geographic distance within ancestries is larger in the continental dataset (AFR/EUR/EAS), the average error is lower, because the method is able to properly discriminate between the three widely separated, divergent ancestries. The within-Asia dataset has closely related (intra-national) ancestries (two from India and three from China), which are very challenging to discriminate via local ancestry, leading to a higher average error.

**Table 5:** Average absolute error (in degrees) of a XGMix model (with window size of 500 SNPs and window smoothing size of 101) on both datasets.

| **Dataset** | **Smoothing** | **Latitude error** | **Longitude error** |
| --- | --- | --- | --- |
| **Continental**<br>**(AFR-EUR-EAS)** | **No** | 4.48 | 26.25 |
|  | **Yes** | 1.18 | 8.46 |
| **Asian**<br>**(10 Classes)** | **No** | 9.14 | 15.01 |
|  | **Yes** | 8.18 | 8.63 |

## References

[1] A. B. Popejoy and S. M. Fullerton, “Genomics is failing on diversity,” Nature News, vol. 538, no. 7624, pp. 161–164, October 2016.

[2] J. Z. Li et al., “Worldwide human relationships inferred from genome-wide patterns of variation,” Science, vol. 319, no. 5866, pp. 1100–1104, February 2008.

[3] M. DeGiorgio, M. Jakobsson, and N. A. Rosenberg, “Out of Africa: modern human origins special feature: explaining worldwide patterns of human genetic variation using a coalescent-based serial founder model of migration outward from Africa.,” Proceedings of the National Academy of Sciences of the United States of America, vol. 106, no. 38, pp. 16057–16062, September 2009.

[4] A. R. Martin et al., “Clinical use of current polygenic risk scores may exacerbate health disparities,” Nature Genetics, vol. 51, no. 4, pp. 584–591, Apr. 2019.

[5] F. Rajabli et al., “Ancestral origin of ApoE *ε*4 Alzheimer disease risk in Puerto Rican and African American populations,” PLoS Genetics, vol. 14, no. 12, pp. e1007791, Dec. 2018.

[6] G. Sirugo, S. M. Williams, and S. A. Tishkoff, “The Missing Diversity in Human Genetic Studies.,” Cell, vol. 177, no. 1, pp. 26–31, Mar. 2019.

[7] A. R. Martin et al., “An unexpectedly complex architecture for skin pigmentation in africans,” Cell, vol. 171, no. 6, pp. 1340–1353, November 2017.

[8] S. Sankararaman, S. Sridhar, G. Kimmel, and E. Halperin, “Estimating local ancestry in admixed populations,” The American Journal of Human Genetics, vol. 82, no. 2, pp. 290–303, February 2008.

[9] B. K. Maples, S. Gravel, E. E. Kenny, and C. D. Bustamante, “RFMix: a discriminative modeling approach for rapid and robust local-ancestry inference,” The American Journal of Human Genetics, vol. 93, no. 2, pp. 278–288, August 2013.

[10] T. Chen and C. Guestrin, “XGBoost: A scalable tree boosting system,” Proceedings of the ACM SIGKDD International Conference on Knowledge Discovery and Data Mining, p. 785–794, 2016, San Francisco, CA.

[11] H. Tang, M. Coram, P. Wang, X. Zhu,, and N. Risch, “Reconstructing genetic ancestry blocks in admixed individuals,” American Journal of Human Genetics, vol. 79, pp. 1–12, May 2006.

[12] A. Sundquist, E. Fratkin, C. B. Do, and S. Batzoglou, “Effect of genetic divergence in identifying ancestral origin using HAPAA,” Genome research, vol. 18, pp. 676–682, April 2008.

[13] A. L. Price et al., “Sensitive Detection of Chromosomal Segments of Distinct Ancestry in Admixed Populations,” PLoS Genetics, vol. 5, no. 6, pp. 1–18, June 2009.

[14] E. Y. Durand, C. B. Do, J. L. Mountain, and J. M. Macpherson, “Ancestry Composition: A Novel, Efficient Pipeline for Ancestry Deconvolution,” bioRxiv, October 2014.

[15] D. Mas Montserrat, C. Bustamante, and A. Ioannidis, “LAI-Net: Local-Ancestry Inference With Neural Networks,” Proceedings of the IEEE International Conference on Acoustics, Speech and Signal Processing, May 2020, Barcelona, Spain.

[16] D. Mas Montserrat, C. Bustamante, and A. Ioannidis, “Class-Conditional VAE-GAN for Local-Ancestry Simulation,” Machine Learning in Computational Biology, December 2019, Vancouver, Canada.

[17] Y. Freund and R. E. Schapire, “A desicion-theoretic generalization of on-line learning and an application to boosting,” Computational Learning Theory, pp. 23–37, 1995.

[18] “http://www.kaggle.com/,”.

[19] 1000 Genomes Project Consortium and others, “A global reference for human genetic variation,” Nature, vol. 526, no. 7571, pp. 68, 2015.

[20] D. Scott, Multivariate Density Estimation: Theory, Practice, and Visualization, A Wiley-interscience publication. Wiley, 1992.

